# Maternal–fetal inflammation affects *Cdh1*/E-cadherin epigenetic regulation and craniofacial development

**DOI:** 10.1101/2025.09.27.676141

**Authors:** Diogo Nani, Lucas Alvizi, Heloisa Maria de Siqueira Bueno, Ellen Cristina Miranda Lacerda, Chao Yun Irene Yan, Maria Rita Passos-Bueno

## Abstract

Cleft lip with or without palate (CLP) is a multifactorial trait associated with environmental exposures such as smoking, alcohol consumption, and pro-inflammatory conditions during early pregnancy, as well as with both common and rare genetic variants. Our group identified *CDH1* loss-of-function variants segregating in families with CLP, with incomplete penetrance linked to promoter hypermethylation. Previously, we demonstrated that gene-environment interactions driven by pro-inflammatory factors can induce this methylation, downregulate E-cadherin expression, and impair neural crest migration, thereby contributing to the etiology of CLP. However, the embryonic consequences of the two-hit CLP model (*CDH1* haploinsufficiency combined with pro-inflammatory insult) have not yet been explored in mammals. Here, under single dose of floxed-*Cdh1*, we investigated the pro-inflammatory response at the maternal-fetal interface, its impact on cytokine and *Cdh1* expression along the anteroposterior axis, the associated epigenetic landscape in the embryonic head, and the effect on cranial structures. We found that pro-inflammatory activation differentially signals to the embryo between anterior and posterior regions, impairing *Cdh1*/E-cadherin expression in NC cells of the head, accompanied by *Cdh1* promoter hypermethylation and other differentially methylated genes involved in cell-junction maintenance. Our findings support a model in which maternal pro-inflammatory responses act as environmental factors that repress NC *CDH1* expression through epigenetic mechanisms, contributing to CLP development by downregulating E-cadherin and potentially compromising overall epithelial integrity. Animal Ethics Committee (CEUA IB/USP) number approval: 394/2022

## Introduction

Cleft lip with or without cleft palate (CLP) is a multifactorial trait considered to be the most prevalent craniofacial malformation worldwide, affecting in average 1:1,000 live-births(1–4), with significant financial and psychosocial impacts(5–8). Aside from suffering impairments in breathing and sleeping(9), swallowing(10) and vocalization(11), CLP represents a significant cost to health systems. A Canadian study observed patients from 1999 to 2021, and estimated the average cost for CLP care at U$73.398,00(12). Also, a study from the Netherlands has estimated the average cost for treatment, between 0 and 24 years of age, as €40.859,00(8). In the state of Paraná, Brazil, it was estimated that treatment complementary to reconstruction surgeries is R$ 18.992,98 (13).

Common and rare genetic variants in genes of cell surface adhesion and mobility molecules, growth and transcription factors, nutrient metabolism enzymes and immune system components have been associated with CLP(14,15). Single nucleotide polymorphisms (SNPs) in genes of morphogens and transcription factors, such as *WNT3*, *WNT9B*, *WNT10A*, *AXIN2* e *MSX1*, are associated with predisposition to CLP with small impact, but significant cumulative effects(16–18). On the other hand, harbouring a single rare variant that affects central genes in the canonic Wnt/β-catenin pathway, such as *CTNNB1*, *AXIN2*, *GSK3B* and *LRP6* confer high risk for CLP(16). Modifications in the proliferation and migration of neural crest (NC), caused by alterations in the cellular cadherin complex, such as the ones related to Epithelial-to-Mesenchymal Transition (EMT), are related to the etiology of craniofacial abnormalities(14,19–25). Some studies have pointed to the role of *CDH1* variants in CLP(23,26,27). *CDH1* encodes for the epithelial cadherin (E-cadherin), which plays an important role in cell adhesion and signalling involved in morphogenesis processes(26–33). Responsible for adherens junctions in cell-cell contact, E-cadherins are key structural factors for epithelial integrity so that null *CDH1* mutant embryos are unviable due to failure in trophectoderm development(34,35). E-cadherin also plays an important signaling role in combination with β-catenin (*CTNNB1*) and p120ctn (*CTNND1*) in the Wnt pathway(36). Disturbances in this function can directly affect migration, polarization and organization of cells during craniofacial development. Evidence in mouse models have shown that alterations in the Wnt/β-catenin pathway, including deletions in *Cdh1*, *Ctnnb1* and *Ctnnd1* result in facial and tooth malformation(36–38). Specifically on NC ontogeny, the EMT-associated E-cadherin to N-cadherin (*CDH2*) switch plays an important role in delamination prior to the migration and differentiation of craniofacial structures(27,39). Therefore, disturbances in NC *CDH1* regulation are thought to affect craniofacial development(4,40,41).

E-cadherin is also known as a tumour suppressor(42) . Mutations in *CDH1* are central to the pathogenesis of Diffuse Gastric and Lobular Breast Cancer Syndrome (DGLBC) (OMIM #137215), an inherited condition that accounts for 1-3% of stomach cancers, which can occur with or without CLP(42,43). *CDH1* pathogenic variants are also linked to autosomal dominant blepharocheilodontic syndrome 1 (OMIM # 119580) (44,45). Loss of E-cadherin drives cells to mesenchymal phenotype, disrupting tissue architecture and promoting invasiveness, a hallmark of DGLBC(43). It also plays a role in other epithelial-derived cancers, such as in oral squamous cell carcinoma, in which E-cadherin expression, as well as podoplanin and vimentin, have been evaluated as a marker of EMT(46). In DGLBC, tumorigenesis typically follows the two-hit hypothesis, wherein a germline *CDH1* pathogenic variant is followed by a second somatic event, such as promoter hypermethylation, loss of heterozygosity, allele-specific silencing, or somatic mutation, leading to E-cadherin inactivation and malignant transformation(46–49). DNA methylation is not limited to cancer progression. It also plays a vital role in developmental disorders. *CDH1* genetic variants(50), and promoter hypermethylation, has been identified in non-syndromic CLP, the latter suggesting a potential epigenetic second hit that contributes to disease penetrance in individuals already carrying germline *CDH1* mutations(49,51).

The “two-hit” hypothesis is a genetic model that explains how certain diseases may arise not from a single genetic mutation, but from the combined effects of two distinct genetic or environmental insults. The first “hit” typically involves an inherited germline mutation that alone may not result in disease but establishes a predisposed state. The second “hit” is usually a somatic mutation or environmental factor occurring later in development, triggering disease manifestation. For instance, the recurrent 16p12.1 microdeletion has been identified as a risk factor for neurodevelopmental disorders, and many individuals carrying this deletion remain asymptomatic unless a second large copy number variation is also present, supporting the two-hit model(52,53). Similarly, in schizophrenia, early disruptions in cell signaling pathways during central nervous system development (first hit) may predispose individuals to later dysfunction when these same pathways are reactivated or stressed during adolescence (second hit)(54). Animal models further support this. Mice with a heterozygous deletion in *Cntnap2* exhibit neurobehavioral impairments only when exposed to early-life stressors, revealing how gene-environment interactions can converge to produce a complex phenotype(55). In hereditary hemorrhagic telangiectasia, vascular malformations, including craniofacial arteriovenous malformations, have been shown to require biallelic inactivation of disease-related genes like *SMAD4*, consistent with a two-hit mechanism involving both inherited and somatic mutations(56). Craniofacial disorders such as craniofacial microsomia further illustrate this. Pathogenic variants in *FOXI3* were often inherited from unaffected parents, suggesting that additional genetic modifiers or environmental exposures may be required to fully express the phenotype(57). The interplay between genetic mutations and environmental insults, such as ethanol, also exemplifies a two-hit scenario. Ethanol-induced craniofacial defects in *Bmp* mutant zebrafish were attributed to increased apoptosis in cranial NC, which are essential for craniofacial morphogenesis(58). These findings underscore the importance of NC survival and signaling interactions with surrounding tissues, which are vulnerable to disruption by concurrent genetic mutations and environmental insults, ultimately driving the spectrum of craniofacial anomalies. Therefore, gene-environmental interactions involved in cell migration and differentiation affect craniofacial cartilage and bone development(15,59–61).

Such gene-environmental interactions such as intrauterine exposure to maternal cytokines, transferred via placental circulation, can induce fetal development modifications of clinical relevance(62–64). Pro-inflammatory activation has been related to impairments involved in orofacial clefts(65–70) with entangled epigenetic regulation(71), and studies have pointed to *CDH1* hypermethylation/E-cadherin downregulation owing to immune activation(72–77). Our group has previously identified heterozygous *CDH1* loss of function variants segregating in families of CLP patients, and associated with incomplete penetrance, estimated as ∼50%(78). It was further demonstrated that CLP penetrance in *CDH1*^+/-^ individuals was positively correlated with *CDH1* promoter hypermethylation in white-blood cells(79). We therefore hypothesized that immune activation and *CDH1* loss of function could synergise in the CLP etiology. Using human induced Pluripotent Stem Cell (hiPSC)-derived NC cells, *Xenopus* and mouse embryos, we previously showed evidence of a CLP two-hit model based on *CDH1* heterozygous loss of function and pro-inflammatory response, as genetic and environmental hits, respectively(51). However, we did not explore the maternal-fetal communication or further embryonic outcomes in the anteroposterior axis Here, we sought to investigate maternal, placental and embryonic immune responses and the repercussions on *Cdh1* expression and epigenetic landscape in the NC of mouse embryos, as well as the craniofacial development. For pro-inflammatory activation, bacterial lipopolysaccharide (LPS) was administered into adult pregnant mice, and the *Wnt1*-Cre *Cdh1*-LoxP system was used to obtain semi-floxed embryos, defined as embryos harboring one floxed *Cdh1* allele in the NC lineage.

## Results

### Maternal exposure to LPS induces systemic and placental pro-inflammatory response

In order to investigate the pro-inflammatory activation as an environmental factor of embryonic epigenetic regulation, intraperitoneal LPS or sterile vehicle control were injected in pregnant females at embryonic stage E8.5. Whole maternal blood and placental tissue samples were collected 2, 4 and 24 hours after the LPS injection (Fig. 1a), as specified in Methods. To get insight into the maternal-fetal interface, we first analysed whole maternal blood Interleukin-1β (*Il-1β*), Interleukin-6 (*Il-6*) and Tumour necrosis factor alpha (*Tnf-α*) mRNA expression at 2, 6 and 24 hours after LPS administration. Two hours from LPS injection, the relative expression of *Il-6* (p = 0.0571) (Fig. 1b) tended to be increased up to 10^4^ times, followed by all three cytokines significant expression augmentation 6 hours after the immune activation induction (Fig. 1c). No statistically relevant differences were observed at 2 or 24 hours points (Fig. 1b, 1d). Therefore, in the systemic expression profile, all three cytokines were significantly transcriptionally upregulated 6 hours after LPS exposure in maternal blood (Fig. 1e). Similarly, we found the same cytokines upregulated in placental tissue at the same 6 hours point (Fig. 1f). Confirming the transcriptional data, protein levels of IL-1β, IL-6 and TNF-α in both maternal serum and placental tissue were evaluated. Except for maternal serum TNF-α (p = 0.3429), all cytokines were found to be at significantly higher concentration in individuals exposed to LPS (Fig. 1g, 1h). Taken together, these results show that the treatment drives significant maternal systemic and placental immune activation detectable within 6 hours.

**Figure 1.**
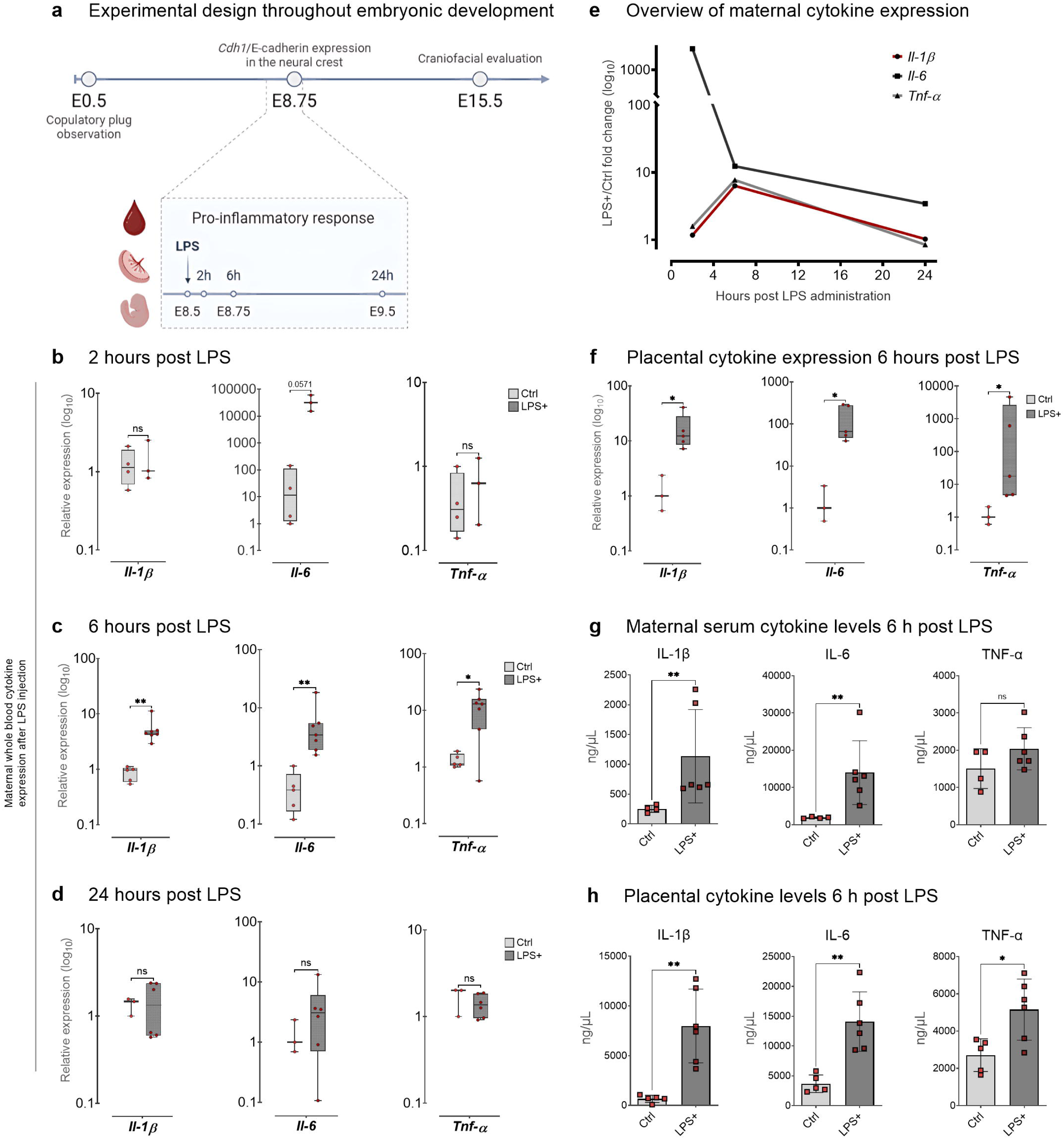
LPS activates systemic and placental pro-inflammatory cytokine expression. **a** Experimental design throughout embryonic development. The copulatory plug marked stage E0.5; LPS was administered at E8.5; pro-inflammatory response was assessed 2, 6 and 24 hours later. Maternal whole blood cytokine expression post LPS injection: **b** 2 hours post LPS injection: no statistically significant differences observed. **c** 6 hours post LPS: *Il-1β* (p = 0.0025), *Il-6* (p = 0.0025), and *Tnf-α* (p = 0.0480) were upregulated in the LPS group **d** 24 hours post LPS: no statistically significant differences observed. **e** Overview of maternal cytokine expression shows a peak at 6 h post LPS, expressed as LPS/control mRNA fold change. **f** Pro-inflammatory cytokine expression in the placenta 6 hours post LPS injection (n(Ctrl) = 5, n(LPS) = 6): *Il-1β* (p =0.03571), *Il-6* (p = 0.0357) and *Tnf-α* (p = 0.0480) were upregulated in LPS-treated individuals. **g** Maternal serum cytokine levels 6 h post LPS (n(Ctrl) = 4, n(LPS) = 6): IL-1β (p = 0.0095) and IL-6 (p = 0.0095) were significantly increased, with a trend for TNF-α. **h** Placental cytokine levels 6 h post LPS (n(Ctrl) = 5, n(LPS) = 6): IL-1β (p = 0.0043), IL-6 (p = 0.0043), and TNF-α (p = 0.0303) were significantly increased. Boxplot whiskers represent minimum to maximum values; in bar plots, lines indicate mean ± standard deviation.

### Impact of maternal–fetal pro-inflammatory activation on embryonic development

We sought to understand how the maternally administered insult activates the embryonic pro-inflammatory response along the anteroposterior axis, its morphological consequences on semi-floxed (*Wnt1*-Cre^+^ *Cdh1*^flox/+^) embryos, 6 hours after LPS maternal injection. LPS-treated embryos exhibited overall loose epithelial morphology (Fig. 2a), but no differences in the analysed morphometric parameters. The pharyngeal arch lateral area (PALA), crown–rump length (CRL), and abdominal antero–posterior diameter (APD) were compared (Fig. 2b). No alteration was observed in any of these features, nor were there differences in PALA relative to CRL or APD according to LPS treatment (Fig. 2c). We next investigated how the maternal pro-inflammatory activation relays to the embryo, and whether this process is related to the regulation of *Cdh1* in the context of cadherin-switch. *Il-1β*, *Il-6*, *Tnf-α* and Nuclear factor kappa B (*Nf-kB*) were evaluated in the anterior and posterior embryonic regions. *Il-6* (p = 0.0087) and *Tnf-α* were observed to be significantly (p= 0.0173) upregulated in the head, and *Il-1β* and *Nf-kB* tended to upregulation (p = 0.0519 and p = 0.0519) (Fig. 2d). Yet no differences in the expression of the same four cytokines were observed in the truncal region (Fig. 2e), suggesting that maternal immune activation affects the embryonic portions of head and trunk differently. The relative expression of genes related to the cadherin-switch and EMT (*Cdh1*, *Cdh2*, and *Zeb1*) was analyzed in the head and trunk regions. *Zeb1* is a key transcription factor controlling EMT and neural crest specification. *Cdh1* was significantly downregulated (p = 0.0088) in the head, whereas the upregulation in *Cdh2* (p = 0.1225) did not reach statistical significance and neither *Zic1* showed to be differentially expressed between the LPS and control groups (Fig. 2f). In the trunk, no differences were observed in the expression of any of the three markers (Fig. 2g). These results indicate that the pro-inflammatory response in the proposed model is more prominent in the embryo head than in the trunk. Furthermore, this anterior-predominant response is accompanied by differential *Cdh1* expression, yet has no evident effect on the size of the first pharyngeal arch within the first few hours of inflammatory response.

**Figure 2.**
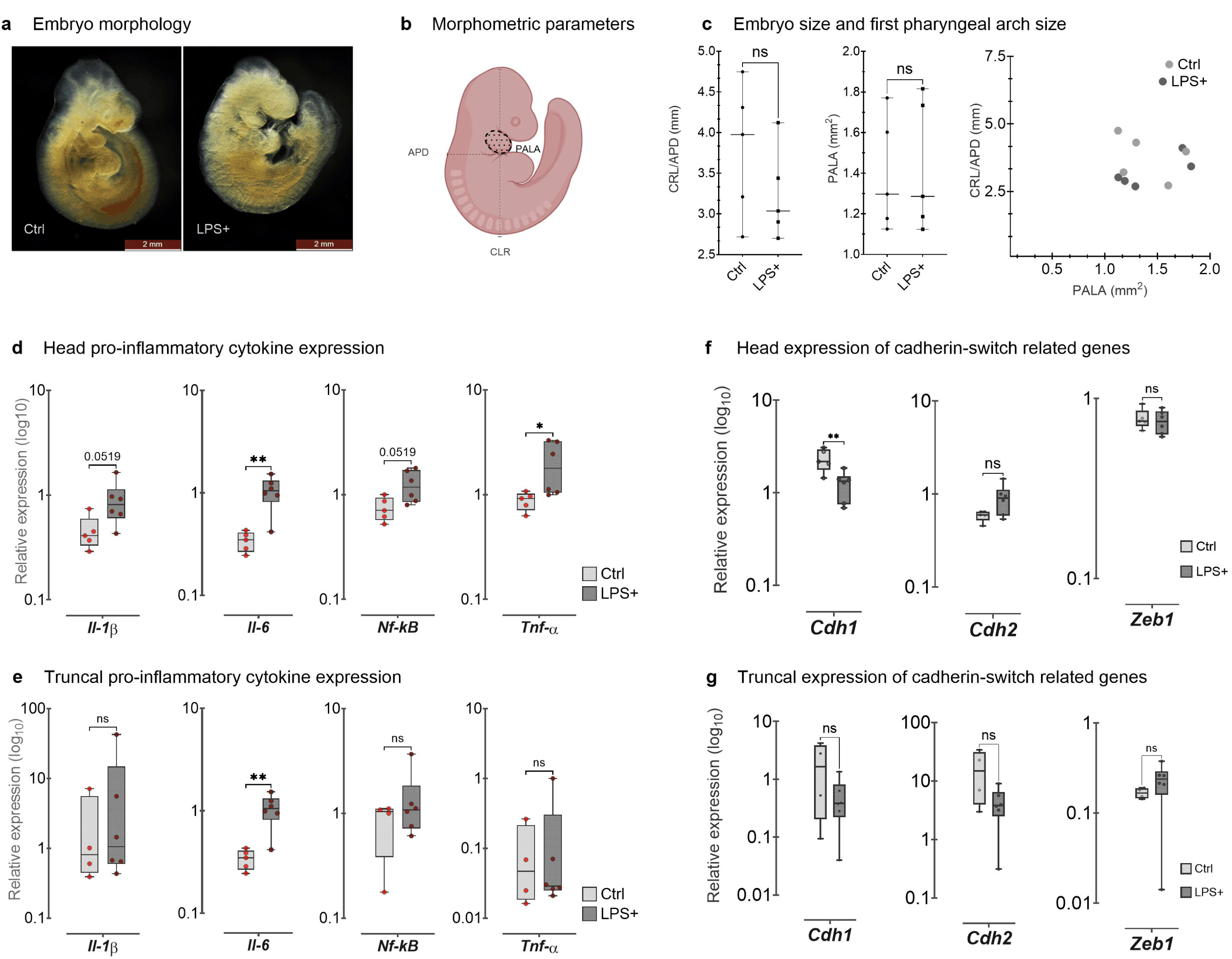
Maternal pro-inflammatory activation induces embryonic axial differences in cytokine and *Cdh1* expression. **a** Embryo morphology under light microscopy shows looser epithelial structure in LPS-treated individuals. **b** Morphometric parameters: first pharyngeal arch lateral area (PALA, black hatched circle), crown–rump length (CRL, grey vertical line), and abdominal antero-posterior diameter (APD, grey horizontal line). **c** Quantification of embryo and first pharyngeal arch size: no differences between LPS and control groups in CRL/APD ratio or PALA. **d** Head pro-inflammatory cytokine expression: *Il-6* (p = 0.0087) and *Tnf-α* (p = 0.0173) were significantly upregulated; *Il-1β* and *Nf-κB* showed a trend towards upregulation. **e** Trunk pro-inflammatory cytokine expression: no significant differences observed. **f** Head expression of cadherin-switch–related genes: *Cdh1* was significantly downregulated (p = 0.0087), while *Cdh2* and *Zeb1* showed no significant differences. **g** Trunk expression of cadherin-switch–related genes: no significant differences in *Cdh1*, *Cdh2*, or *Zeb1*. Lines indicate median and 95% CI; boxplot whiskers represent minimum to maximum values.

Following the transcriptional analysis, we focused on the head region to determine whether E-cadherin downregulation occurs specifically at the protein level in NC cells. To address this, we measured E-cadherin protein abundance in Sox10^+^ cells. We observed a significant decrease in the ratio of E-cadherin^+^ to E-cadherin^-^ cells within the Sox10^+^ population of the LPS group (p = 0.0043); this difference was not statistically significant in the overall cell population (Fig. 3a).The proportion of Sox10+ cells was similar between the LPS and control groups (46.7% and 54.9%, respectively). However, the percentage of E-cadherin-cells was higher in the LPS group (33.1%) compared to the control (28.3%), indicating a treatment-specific effect. To rule out the possibility that the treatment altered neuroepithelial identity, we characterized the expression of key lineage markers: NC markers (*Ap2b*, *Sox10*), neural markers (*Notch1*, *Pax6*), and epithelial markers (*Cldn6*, *Krt19*). No significant differences in mRNA expression were observed between groups (Fig. 3b). We next investigated whether *Cdh1* downregulation was mediated by epigenetic mechanisms.

**Figure 3.**
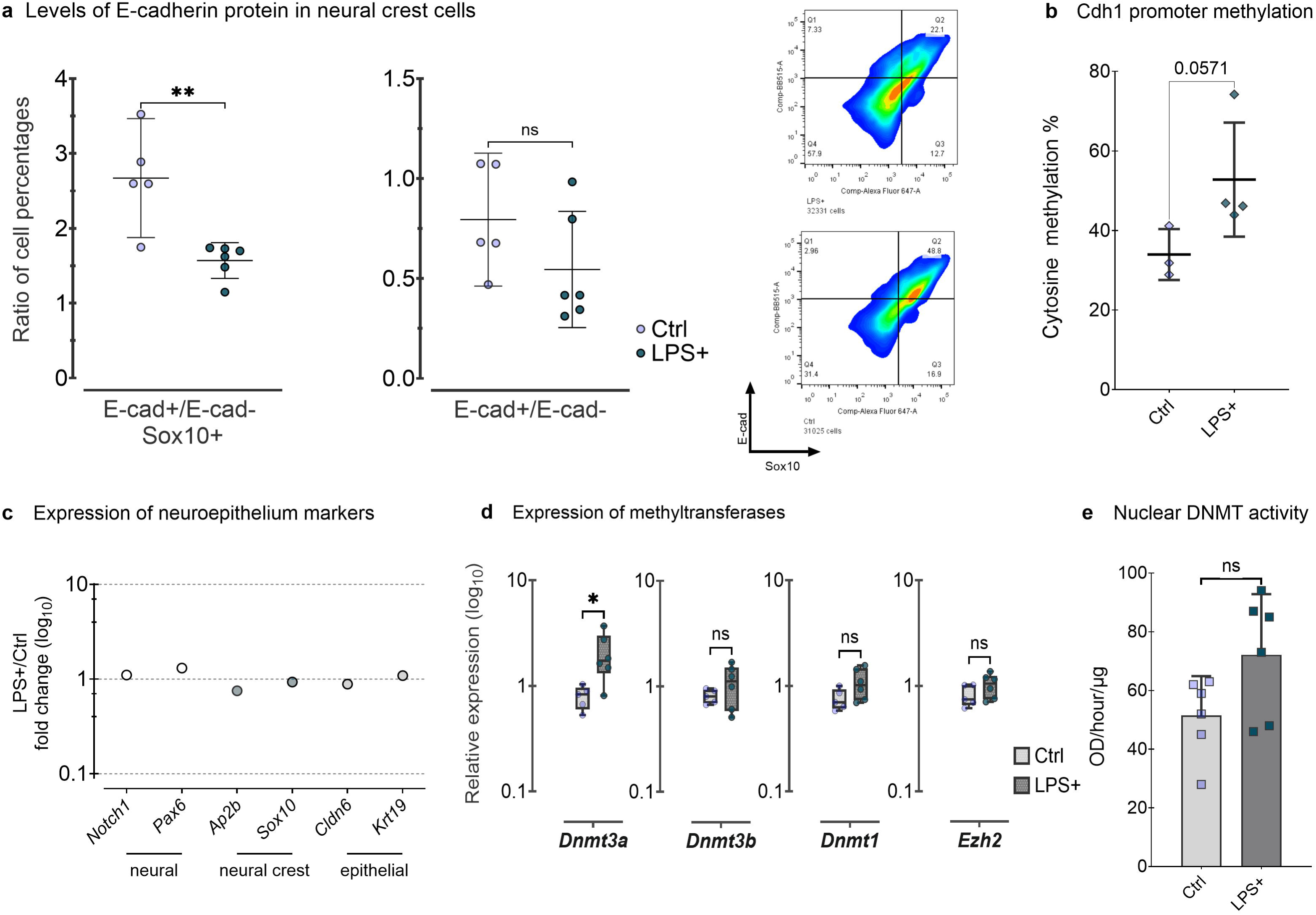
Pro-inflammatory response in the embryo head epigenetically downregulates neural crest *Cdh1*/E-cadherin. **a** E-cadherin protein levels in neural crest and head region cells: the E-cadherin+/E-cadherin− ratio was significantly decreased in Sox10+ cells (p = 0.0043), while no significant difference was observed for all analysed cells. In the heat map, E-cadherin and Sox10 are displayed on the x- and y-axes, respectively. **b** *Cdh1* promoter methylation tends to be higher in the LPS group. **c** Expression of neuroepithelium markers: no significant changes in LPS/control fold change. **d** Expression of methyltransferases: only *Dnmt3a* was significantly upregulated (p = 0.0269). **e** Nuclear DNMT activity in LPS-treated embryos was not significantly different from controls. Lines indicate median and 95% CI; boxplot whiskers represent minimum to maximum values; column lines indicate mean ± SD.

We evaluated the mRNA expression of DNA methyltransferases (*Dnmt1*, *Dnmt3a*, *Dnmt3b*) and the histone methyltransferase *Ezh2*. Only *Dnmt3a* was transcriptionally upregulated (p = 0.0303) in LPS-treated embryos; no differences were observed for *Dnmt1*, *Dnmt3b*, or *Ezh2* (Fig. 3c). To investigate genome-wide DNA methylation changes, including those affecting Cdh1, we performed Reduced Representation Bisulfite Sequencing (RRBS). On a genome-wide scale, the LPS group exhibited 37.2% methylation in the CpG context, 3.4% in CHG (H = A, T, or C), and 2.7% in CHH, whereas the control group showed 33.8%, 3.1%, and 2.5%, respectively. The distribution of methylated CpGs revealed that 0.52% were located within CpG islands, 3.30% in shores, and 96.18% in other genomic regions.

Differential methylation analysis identified 1% hypomethylated and 3% hypermethylated regions (DMRs), with their chromosomal distribution shown in Figure 5a. No significant difference in global methylation levels was detected between groups (t-test, p = 0.1091). Gene Ontology (GO) analysis of DMRs revealed enrichment for cellular component maintenance and cell junction maintenance and (Table 1). We validated *Cdh1* promoter hypermethylation by bisulfite sequencing, which showed a strong tendency towards an increased cytosine methylation (p = 0.0571) in the LPS group (n = 4) compared to the control (n = 3). These results demonstrate that the maternal inflammatory insult alters the epigenetic landscape of *Cdh1* and affects methylation genome-wide. Furthermore, the downregulation of E-cadherin protein in neural crest cells (Fig. 3a), coupled with unchanged lineage marker expression (Fig. 3b), indicates that E-cadherin loss is a targeted event that occurs without broader changes to neuroepithelial cell identity, driven by epigenetic modifications modulated by the pro-inflammatory activation.

**Table 1.**
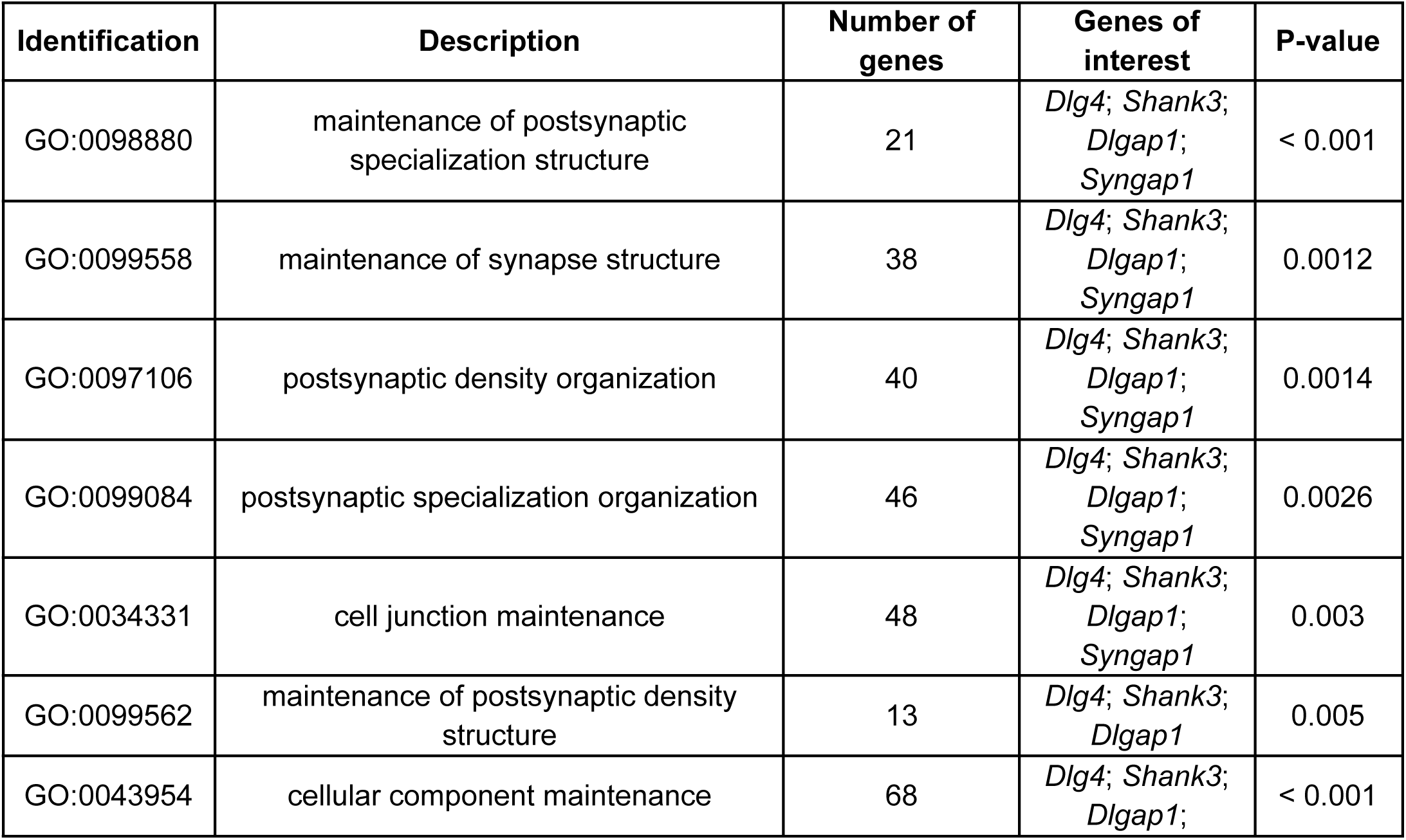
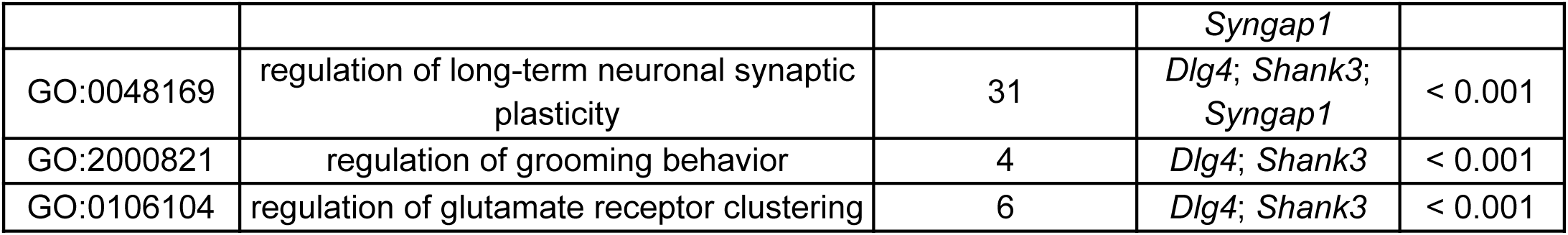
Top 10 enriched biological processes (GO terms) identified by WebGestalt for differentially methylated regions (DMRs) in head tissue, in LPS-treated Cdh1 semi-floxed embryos relative to controls.

Lastly, we asked whether the combination of inflammatory insult and *Cdh1* haploinsufficiency could drive craniofacial malformations. To test this, pregnant Cre-LoxP and wild-type females were injected with LPS at E8.5, and embryos were harvested at E15.5 for morphological assessment. Semi-floxed embryos frequently exhibited neural tube closure defects (83% penetrance, 5/6), whereas this phenotype occurred at lower frequency in *Wnt1*-Cre^+^ embryos (33% penetrance, 1/3) and was absent in wild-type controls (n=4) (Fig. 4a). In addition, semi-floxed embryos displayed severe craniofacial abnormalities, including neurocranial defects (83% penetrance, 5/6), cleft palate (50% penetrance, 3/6), and wider lip gaps (50% penetrance, 3/6) (Fig. 4a,b). Notably, morphometric analysis revealed that only semi-floxed embryos exhibited significantly wider lip gaps compared with wild-type (p = 0.0069), while *Wnt1*-Cre^+^ showed intermediate lip gap sizes (Fig. 4b). Although *Wnt1*-Cre^+^ littermates also presented wider lip gaps relative to wild-type, this difference did not reach statistical significance (Fig. 4b). Also, semi-floxed embryos were significantly smaller than wild-type, as indicated by a reduced CRL/APD ratio (p = 0.0197), while *Wnt1*-Cre^+^ embryos showed intermediate values (Fig. 4c). These results indicate that the Cre driver itself may contribute to the craniofacial phenotype, whereas maternal pro-inflammatory exposure alone is insufficient to reproduce the observed defects. Taken together, our findings suggest that the combination of a genetic factor, mediated by single dose of *Cdh1* Cre-LoxP silencing, and an environmental factor, driven by maternal pro-inflammatory insult, can synergistically lead to craniofacial impairments.

**Figure 4.**
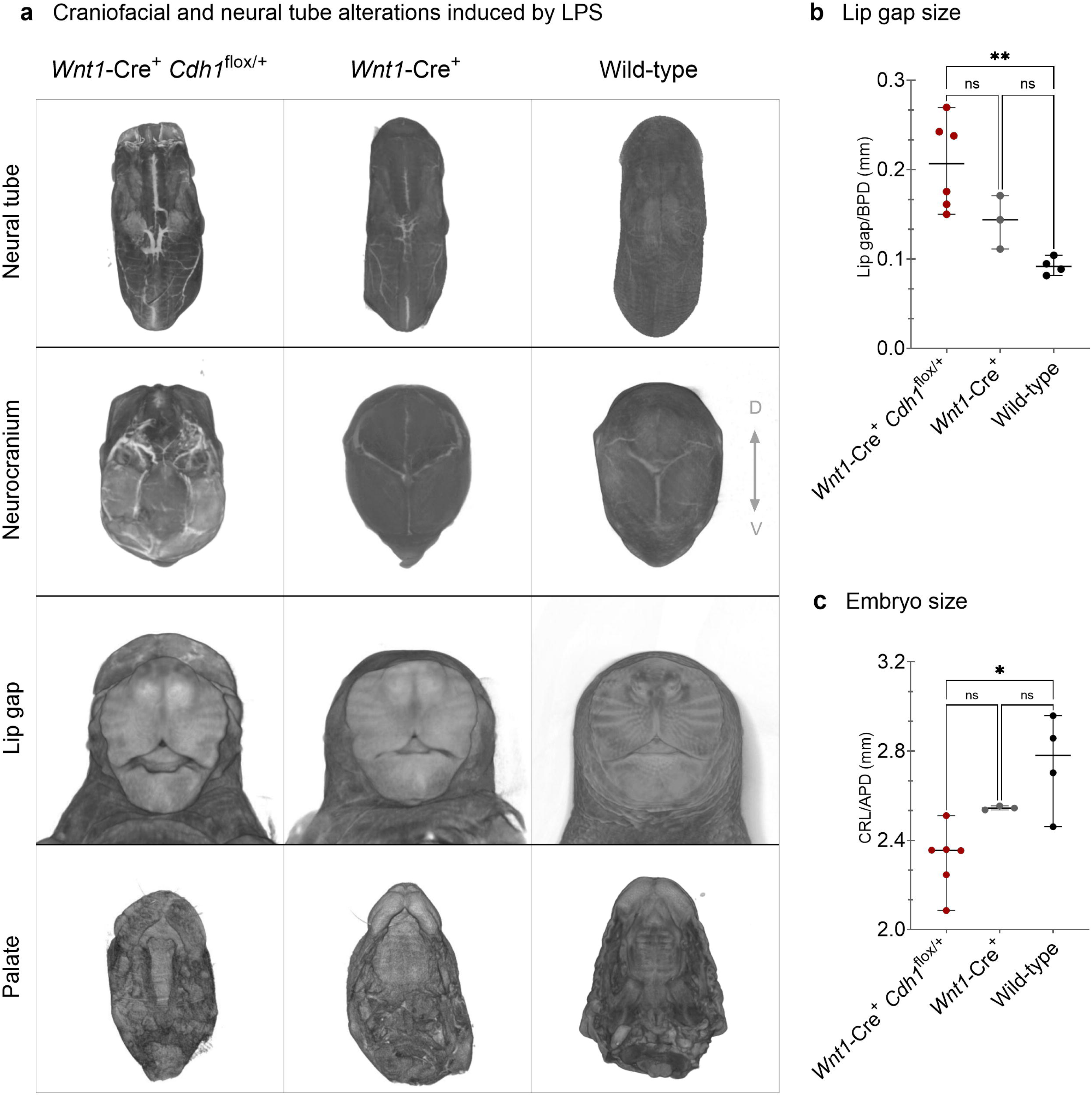
*Cdh1* single dose silencing, in combination with pro-inflammatory insult, drives craniofacial and neural tube malformations. **a** Craniofacial and neural tube defects induced by LPS in *Wnt1*-Cre^+^ *Cdh1*^flox/+^ embryos (n = 6) include neural tube enclosure defects (83% penetrance), neurocranium malformations (83%), increased lip gap size (50%), and cleft palate (50%). In *Wnt1*-Cre+ embryos, neural tube enclosure defects were observed with 33% penetrance. **b** Lip gap size showed to be greatly increased in *Wnt1*-Cre^+^ *Cdh1*^flox/+^ in comparison to the wild-type. **c** Embryo size showed to be decreased in *Wnt1*-Cre^+^ *Cdh1*^flox/+^ in comparison to the wild-type. Lines indicate median and 95% CI.

## Discussion

Our study demonstrates that maternal immune response affects the cranial NC and derived structures, and replicates the *Cdh1* downregulation driven by pro-inflammatory epigenetic regulation in the placental model. By exploring the transcriptional activity of pro-inflammatory cytokines in the maternal, placental and embryonic immune response, we demonstrated its systemic response throughout time in pregnant females as well as the embryonic consequences in the anteroposterior axis. Considering the CLP families carrying pathogenic *CDH1* variants that provided the initial evidence for this line of research, placental transmission of immune activation, and its relay to the embryo as an environmental factor that downregulates *Cdh1*/E-cadherin in cranial NC cells through broad epigenetic modulation of cell-junction pathways, represents a key step in supporting our hypothesis.

Through comparative mRNA expression and cytokine quantification, we observed the peak of maternal systemic pro-inflammatory activation 6 hours after LPS induction. In most studies involving LPS treatment in mice, samples are harvested within the first 4 hours, which is a well-established time to investigate inflammatory outcomes(80–82), with a few ones also interested in a longer observation as we were(83,84). Intrauterine embryogenesis is thought to require maternal immunological adaptation as the fetus, a genetically distinct individual, must be supported to grow inside the pregnant female body(85) and a few studies have focused on the characterization of those adjustments. Zöllner and collaborators reported that, in murine models of sepsis, pregnancy at E16, a stage equivalent to the third trimester of human gestation, induces an increase only in the anti-inflammatory cytokine IL-4 in maternal serum, whereas the mRNA expression of IL-10 and the pro-inflammatory cytokines IL-1β, IL-6, and TNF-α is upregulated in myocardial tissue(84). They harvested samples 6 and 12 hours after pro-inflammatory induction, and used 10 μg of LPS regardless of the maternal mass, whilst we administered it according to the animal weight (i.e. 2 μg of LPS for a 20g pregnant mouse, or 100 μg/Kg). On the other hand, Vizi and collaborators reported that also in the third semester, using 10 or 50 mg/kg of LPS and sampling after 1.5 or 4 hours, the levels of IL-6, IFN-γ and TNF-α a are increased in maternal plasma, and the anti-inflammatory IL-10 is decreased(82). They also showed that the rise in circulating TNF-α already occurs at 7–8 days of murine gestation, a stage equivalent to the first trimester of human pregnancy(82). So the results regarding immune activation changes according to the methodology and also the type and quantity of LPS used can also alter the outcomes (86,87). However, these data show that during pregnancy, the pro-inflammatory activity is settled from a different baseline than non-pregnant individuals.

Pro-inflammatory activation was induced in E8.5 embryos via maternal LPS administration, which is able to cross the placental interface(88) and known to upregulate placental TNF-α, IL-1β and IL-6(89), as well as affect epigenetic inheritance in mice(90). Timing for pro-inflammatory induction was based on the offspring average NC migration and development of the choriovitelline placenta(91–93). Although mouse embryonic development resembles human embryogenesis, there are particular properties in the placentation between the two species that should be taken into account in the interpretation of mouse-model derived evidence(93). They have similar placental composition with differences in timing and mediators of decidualization, as well as tissue organization (94–96). In mice, decidualization begins at E3.5, whereas in humans there is a predecidualization before fertilization(97). By E8, mice develop a primitive placenta that is not formed in human gestation, consisting of the juxtaposition of the yolk sac against maternal tissues and vasculature; at E12.5 the yolk sac is displaced by the definitive placenta followed by villi vascularization and around E14.5, the syncytiotrophoblast directly receives maternal blood(97). In humans, the primitive intervillous space forms around days 8–9 of gestation, and the definitive villous structure of the placenta is established by day 21. The fetoplacental circulation becomes fully functional only between the 10th and 12th weeks, when maternal blood begins to perfuse the intervillous space(93,97).Therefore, the stage at which we administered LPS (E8.5 in mice) corresponds to the late pre-vascular phase of murine placental development, approximately equivalent to the stage preceding villous vascularization in the human placenta, around days 16–18 of gestation(93). Although we present results in a mammalian system, we do not know exactly how the induced inflammation represents a human gestational inflammatory scenario, which can be driven by non infectionary scenarios, such as diabetes and nutrition habits (98,99).

Also using the Cre-LoxP system, we have previously reported that maternally administered LPS diminishes NC migration in a *Cdh1*/E-cadherin dose dependent manner in mouse embryos and also showed that NC *Cdh1* knockout (*Wnt1*-Cre^+^ *Cdh1*^flox/flox^ embryos) develops CLP phenotypes(51). We found that *Cdh1* semi-floxed embryos exposed to pro-inflammatory activation exhibited markedly worsened phenotypes, particularly involving craniofacial sctructures and neural tube development. This pattern not only reveals inherent limitations of the model but also emphasizes the wide-ranging impact of systemic immune activation. We did not explore the baseline *Cdh1* expression of this model in comparison to other lineages; however, wild-type embryos submitted to LPS treatment did not develop any observable craniofacial phenotype. Human NC migration was reported to be impaired due to interferon-β exposure(100), and Zika virus infection was shown to induce NC cytokine production(101). In cell metastasis, IL-6 enhances N-cadherin expression(102), and in CLP genes related to pro-inflammatory maintenance are significantly upregulated in child cleft mucosa(103). As reported in the mouse model, pregnancy also alters the immunological system in humans (104,105). Interestingly, it has been reported that pregnant women of black ancestry exhibit greater TNF-α production during mid-pregnancy and marginally lower IL-1 production at postpartum, in comparison to those with caucasian ancestry (106), and it is known that the frequency of CLP is lower in African populations (107). In this context, further studies with other candidate genes involved in EMT might contribute to a better understanding of its role in CLP etiology.

In the experimental conditions, no differences on neural and non-neural epithelia identity were observed, neither on morphology size and proportions. Also, since no NC markers were found differentially expressed under the pro-inflammatory insult, it is suggested that NC induction remains properly maintained in the 2-hit model, but NC migration is affected by the environmental factor, as previously reported (51). The transcriptional activation of pro-inflammatory cytokines was more prominent in the head than in the trunk, as well as the alteration in *Cdh1* expression and DNMT activity, which could be related to the axial specificity of the impairments. The axial differential sensibility to the maternal immune activation occurs also in accordance with reported ability to reactive oxygen species generation(108), which are biologically connected processes(109). Other authors have demonstrated that the cranial NC cells are also particularly sensitive to apoptosis induced by oxidative stress(110,111). In the mammalian embryonic development, cells from the dorsal anterior ectoderm differentiate into neural plate border cells and then into cranial NC, when they go through EMT and acquire mesenchymal phenotype(112–118). NC cells migrate ventrolaterally to the pharyngeal arches and give rise to the five facial prominences: two maxillary, two mandibular and one frontonasal(118). Maxillary prominences form the upper lip, mandibular form the lower lip and jaw, whereas the frontonasal prominence forms the forehead, the nose and the top of the primitive mouth(119). Therefore, NC delamination and migration are sensitive stages of craniofacial ontogeny dependent on EMT(4,41). The cellular phenotypic transition is tightly regulated and it was shown that E-cadherin to N-cadherin switch is orchestrated by ZEB1 and β-catenin activity in other EMT models(73,120). In our study, we did not explore the dynamics involving EMT regulation further than embryonic *Zeb1* transcription, which did not show to be affected by maternal immune activation. Moreover, an E-cadherin/NF-κB regulatory loop was shown to regulate E-cadherin transcription in cancer collective cell migration(121), which is consistent with our proposed gene-environment interaction in the two-hit CLP model.

Together, Sanger sequencing and RRBS results indicate *Cdh1* promoter differential methylation under pro-inflammatory activation, as well as for genes related to cell-junction maintenance. *Dnmt3a* was shown to be upregulated from the four methylating enzymes analysed by RT-qPCR. Noteworthy, *Dnmt3b* was shown to be dispensable for mouse embryonic development elsewhere(128). The disruption of NC cells DNA methylation was shown to drive mouse orofacial clefts(129) and inflammatory response has been related to DNMT3A activity(130,131). In mouse embryonic stem cells, DNMT1 and 3A/B are necessary for CG methylation symmetry across generations of cultured cells(132).

Specifically, DNMT3A/B add methyl groups to recent synthesized DNA strands(131) and DNMT1 maintains DNA methylation during cell division(133). In contrast to our findings, Purkait et al. (2022)(134) showed that pro-inflammatory activation due to *Helicobacter pylori* infection activates *EZH2* and *DNMT1* in the stomach, affecting the expression of *CDH1* among other genes in hereditary diffuse gastric cancer(128). In addition to CpG methylating enzymes, we also analysed the expression of *Ezh2*, that adds methyl groups to histone H3 and was reported by the same group to mediate pro-inflammatory induced epigenetic alterations(135,136). We found enrichment of methylation related to the *Eed*, which encodes a core component of the Polycomb Repressive Complex 2 (PRC2), that plays a pivotal role in maintaining transcriptional repression through H3K27me3 (137). The study by Ying et al. (2019) revealed that EED promoter hypomethylation is significantly associated with colorectal cancer, with tumor tissues exhibiting markedly lower methylation levels compared to normal tissues. In EMT, EED-mediated PRC2 activity can indirectly suppress *CDH1* (E-cadherin) by recruiting EMT transcription factors like *SNAIL* and *ZEB1* to the *CDH1* promoter, thereby facilitating mesenchymal transitions (137). Lastly, *Shank3*, known for its protective effect against inflammation in neurons and associated with autism/Phelan-McDermid syndrome (OMIM #606232) (138,139), which includes craniofacial dysmorphisms (140,141), was found hypomethylated in the experimental group (142).

Considering the E-cadherin structural function in mechanical forces of cellular aggregation, the protein also plays important roles in collective cell migration in a variety of models(121–126), including in orofacial clefts(126). These are fundamental proteins of general epithelia, and inflammatory activation mediated by the placenta is thought to affect the whole embryo. Here, we did not investigate the effect of *Cdh1* loss of function in other tissues, nor investigated other genetic expression alterations induced by immune response in the etiology of CLP. Also, whether the *CDH1* hypomorphism affects NC delamination is still to be investigated, albeit the observed downregulation of Cdh1/E-cadherin is consistent with the reported NC migration impairment(51). Besides acting as an adhesion molecule, E-cadherin also participates in the Wnt pathway by interacting with catenin, which has also been related to CLP(37,78). As a complex condition that affects the most sophisticated human bone and cartilage structure, alterations in cellular cadherin profile due to immune activation, could affect several phenomena involved in craniofacial development—including cellular signalization and morphogenetic movements not necessarily related to the E-cadherin to N-cadherin switch. For example, in chicken embryo NC, pro-inflammatory activation was shown to upregulate both E-cadherin and N-cadherin(127).

## Conclusions

Our findings indicate that maternal pro-inflammatory activation transmitted from pregnant mice to their offspring via placental communication results in defects of NC-derived structures and epigenetic modulation of genes involved in cell junctions. In this model, the inflammatory insult, acting as an environmental factor, predominantly impacts the anterior embryonic region, the site of critical morphogenetic processes for craniofacial development, including NC migration. This mechanism may account for the incomplete penetrance of familial CLP cases harbouring *CDH1* variants, as epigenetic modulation appears to mediate the interplay between genetic susceptibility and pro-inflammatory activation. This study demonstrates how genetic susceptibility and maternal inflammation interact in the etiology of CLP, highlighting *CDH1* as a clinically relevant target for genetic counselling. Importantly, it provides functional evidence that maternal and placental inflammatory signaling in a mammalian model can reshape embryonic development, underscoring translational relevance and possible implications for prenatal care.

## Methods

### Ethics

The maintenance, procedures and experiments of mouse lineages were approved by the Research Ethics Committee from the Biosciences Institute (University of Sao Paulo, Brazil) under the protocol 394/2022. All animal procedures were performed under the Committee’s ethics standards.

### Neural crest–specific Cdh1 conditional heterozygous mice

To generate *Mus musculus* embryos heterozygous for *Cdh1* in neural crest (NC) cells, we crossed *Wnt1*-Cre2 mice (129S4.Cg-E2f1Tg(Wnt1-cre)2Sor/J; Strain #022137, The Jackson Laboratory, USA), which express Cre-recombinase in *Wn1+* cells, with *Cdh1*-LoxP mice (B6.129-Cdh1^tm2Kem/J; Strain #005319, The Jackson Laboratory, USA). When specified, C57BL/6J embryos were used as wild-type controls. In this study, the genotype *Wnt1*-Cre^+^ Cdh1^flox/+^ designates NC *Cdh1* heterozygous embryos, whereas *Wnt1*-Cre^+^ carriers with intact *Cdh1* alleles, considered controls. Embryonic age was estimated as E0.5 based on the detection of a copulatory plug.

### Embryonic morphometric measurements

E8.75 embryo images were acquired under Leica M165 FC light microscope with Leica DFC7000 camera. The pharyngeal arch lateral area (PALA), crown-rump length (CRL) and abdominal antero-posterior diameter (APD), following what was previously described(143). E15.5 craniofacial development was assessed using ultramicrotomography (µCT). To evaluate the lip gap, we calculated the ratio of the upper lip gap to the total upper lip length, and normalized this value to the biparietal diameter (BPD). The resulting values were normalised with respect to both the CRL and the APD, in order to account for overall embryonic size and growth variability. µCT results were obtained with the SkyScan 1176 instrument, using 0.22 aluminium filter with 9 µm resolution, and 3D reconstructed using NRECON v2 software. Images were obtained using the volume rendering software CTvox v3.3.1.0 (Bruker). All measurements were performed on ImageJ software(144).

### Maternal pro-inflammatory activation

For pro-inflammatory activation via maternal immune activation, pregnant mice at E8.5 were injected with 100 µg/Kg LPS from Escherichia coli 026:B6 (Invitrogen™, 00-4976-93) (145). In the control group, pregnant mice were injected with the same volume of sterile vehicle control (1X PBS). Females were placed in experimental or control groups randomly. In order to preserve the effect on pro-inflammatory activation, animals were euthanized with cervical dislocation (^146–148^). Adults and embryos were genotyped by PCR with Platinum Supermix (Invitrogen™) following The Jackson Laboratories recommendations. DNA samples were extracted with NucleoSpin Tissue kit (Macherey-Nagel) and PCR products were separated with electrophoresis in 1,5% agarose gel with GelRed/BFB (1:500) in TBE buffer. Gel images were photo documented with ImageQuant Las 400 mini (GE Healthcare).

### mRNA expression

The expression of NC markers, cytokines, methylating enzymes and cadherins genes were analysed with quantitative Reverse Transcription Polymerase Chain Reaction (qRT-PCR). Samples were individualised in 0.5 ml tubes from properly euthanized animals and dissected in 1X PBS. For anteroposterior analysis, the embryos were transversely sectioned just caudal to the first pharyngeal arch. NucleoSpin TriPrep kit (Macherey-Nagel) was used for RNA extraction, according to recommendations. RNA samples were quantified with spectrophotometer (Epoch, BioTek) and filtered by quality of Abs260/280 = 2 ± 0.1 and Abs260/230 = 2 ± 0.2 ratios (149). cDNA was retro-transcribed with SuperScript™ VILO™ Master Mix (ThermoFisher Scientific) using 1.5 ug of RNA per sample. Fast SYBR Green Master Mix 2X (Applied Biosystems) was used for cDNA amplification with 100nM forward and reverse, following fabricant recommendations. Primer sequences, described in Table 3 were designed in exon-exon junctions using NCBI Primer Blast. Reactions were performed in a QuantStudio 5.0 System (ThermoFisher Scientific) using standard parameters. Relative expression values were calculated as previously reported, using *Actb*, *Gapdh* or *Tbp* as endogenous control selected with NormFinder(150,151).

### Cytokines quantification

We quantified cytokines concentrations in maternal serum and placental protein extractions, from properly euthanized animals, with Enzyme Linked ImmunoSorbent Assay (ELISA) technique. Whole maternal blood was incubated at RT for 30 min to clot and centrifuged for 20 min at 4°C and 16,000xg. Serum containing supernatant was transferred to ice cold tubes and stored at -80°C. Samples were diluted in RIPA Buffer (Thermo Fisher Scientific) (1:50). Pellets were discarded. Placentas were dissected in ice cold 1X PBS. Every 5 mg of placental tissue was transferred to ice cold 1.5 ml tubes containing 300 ul of RIPA Buffer. Samples were homogenised with an ultrasonic sonicator processor (Heat Systems) with 12 cycles of 10s and 30s intervals, with 180W wattage and at 4°C. Next, they were centrifuged at 16,000xg for 20 min at 4°C. Protein containing supernatant was transferred to new ice cold tubes and stored at -80°C. Serum and placental samples were with Pierce™ BCA Protein Assay Kit (Thermo Fisher Scientific). IL1-β, IL-6 and TNF-α levels were measured with arigoPLEX® Mouse Proinflammatory Cytokine Multiplex ELISA Kit (IL1 beta, IFN gamma, TNF alpha, IL6) kit (arigobio). Protein concentrations were estimated with optical density (OD) measured in a spectrophotometer at 450 nm(Epoch, BioTek). Experimental procedures and quantification were performed according to fabricant guidelines.

### Flow cytometry

E-cadherin and Sox10 expressions were evaluated with flow cytometry. Cells were obtained as described by Ward et al. (2023)(^152^). Briefly, dissected head embryos were dissected and tissues were incubated in 500 ul PBS 1x 450 U/ml Collagenase IV 20mM HEPES for 30 min at 37°C and 250 rpm. Cells were separated with 70 μm filters (Corning® Falcon® Cell Strainer) and centrifuge for 5 min at 400xg and 4°C. Single cells were fixed and permeabilized with True-Nuclear™ Transcription Factor Buffer Set (BioLegend), according to manufacturer orientation. The primary antibodies Sox10 (D5V9L) Rabbit mAb #89356 (Cell Signaling) and anti-E Cadherin antibody [M168] ARG41292 (arigobio) were used in 1:100 concentration and the secondary antibodies Anti-rabbit IgG (H+L), F(ab’)2 Fragment (Alexa Fluor® 647 Conjugate) #4414 (Cell Signaling) and Anti-mouse IgG (H+L), F(ab’)2 Fragment (Alexa Fluor® 488 Conjugate) #4408 (Cell Signaling) with 1:500. Analysis was performed using FlowJo™ v10.8 Software (BD Life Sciences) (153).

### DNA methyltransferase activity

Embryonic DNMT3A, DNMT3B and DNMT1 activity was measured with DNMT Activity Quantification Kit *(Colorimetric)* (abcam). Single cells were obtained as described and digestion was verified with flow cytometry, as previously described (152). Supernatant was recovered and cellular dissociation was verified with flow cytometry. Next, NE-PER Nuclear and Cytoplasmic Extraction Reagents kit (Thermo Fisher Scientific) was used for nuclear fractionation and samples were quantified with *Pierce™ BCA Protein Assay Kit* (Thermo Fisher Scientific). Activity was estimated based on OD measurements at 450 nm performed in a spectrophotometer (Epoch, BioTek), according to fabricant guidelines.

### DNA methylation

Reduced representation bisulfite sequencing was used for investigation of diffentialy methylated regions (DMRs). 50 ng of DNA was used for library preparation and downstream sequencing with the Zymo-Seq RRBS Library Kit (Zymo Research). Library quality was assessed with Qubit (Thermo Fisher) and Bioanalyzer (Agilent), and paired-end sequencing was done by the Human Genome and Stem Cell Research Center sequencing service, the using a SP Reagent Kit v1.5 (100 cycles) on a NovaSeq 6000 lanes (Illumina). Quality assessment was performed using FastQC. Reads were mapped to *Mus musculus* NCBI genome (mm39), and methylation calls were extracted with Bismark 0.24.2, using bowtie and methylation extractor modes, respectively (154). Differentially methylated regions (DMRs) were identified with methylkit 3.21 (155), using NCBI RefSeq mm39 genome as reference, 25% as threshold and p < 0.001 as criteria for relevance. Finally, DMRs were analysed in the online platform WebGestalt for biological interpretation; GO were enriched when p < 0.05 (156). Bisulfite Sanger sequencing was used for assessing *Cdh1* promoter methylation with bisulfite-specific primers designed on MethPrimer (157) (Table 2). DNA samples were extracted from the head of E8.75 mouse embryos using NucleoSpin TriPrep kit (Macherey-Nagel) and 200 ng of DNA per individual was converted using BisulFlash DNA Modification Kit (EpigenTek). Amplicons were purified with Sephadex LH-20 (GE Healthcare) and Sanger sequencing was performed at the Human Genome and Stem Cell Research Center. Cytosine peak values in CpGs were called using Sequencher® DNA sequence analysis software (Gene Codes Corporation) (158) and C signals were used to determine the methylation percentage as in 100 * C/(C + T).

**Table 2.**
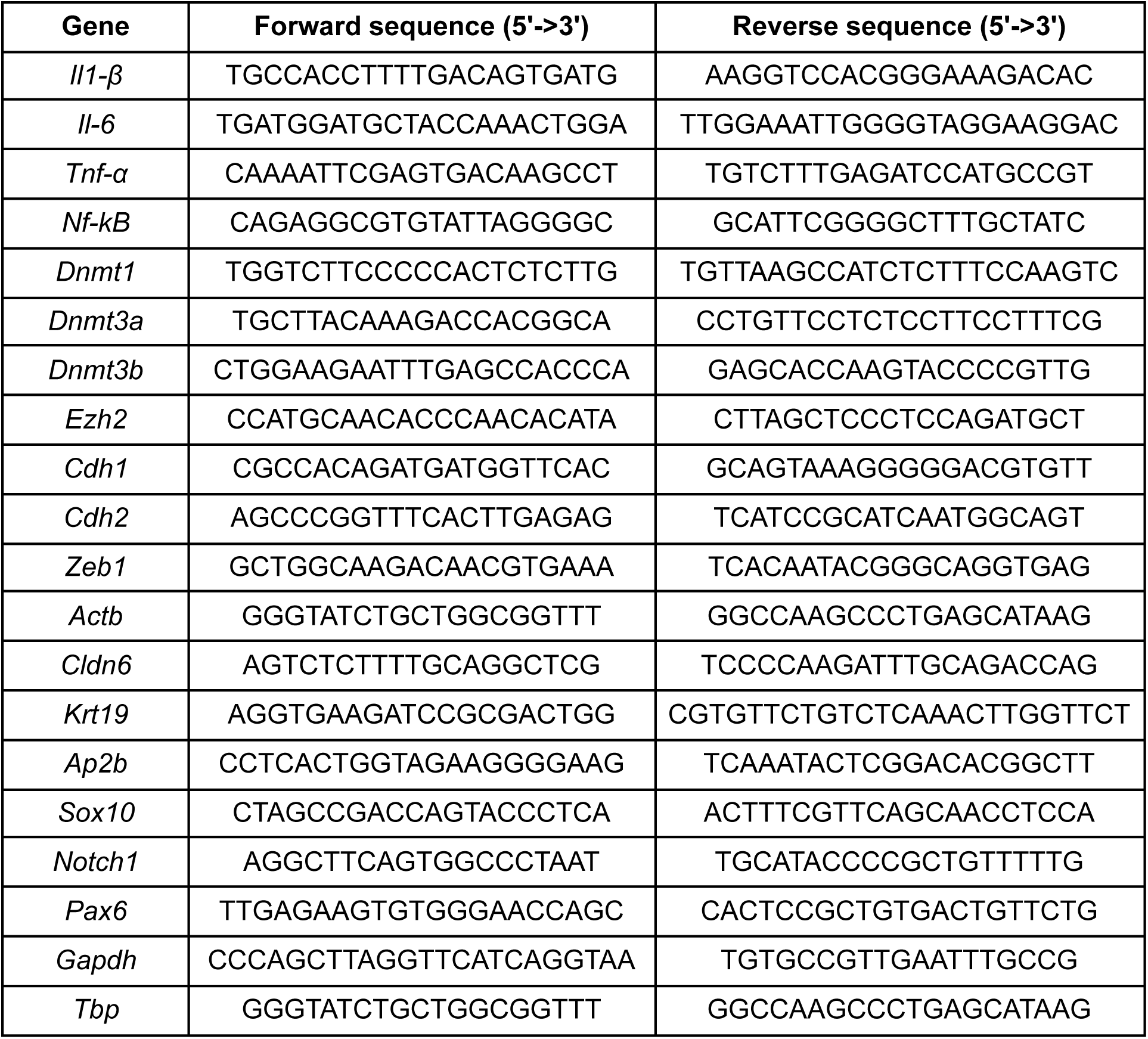
Primer sequences used for qRT-PCR assays. .

**Table 3.**
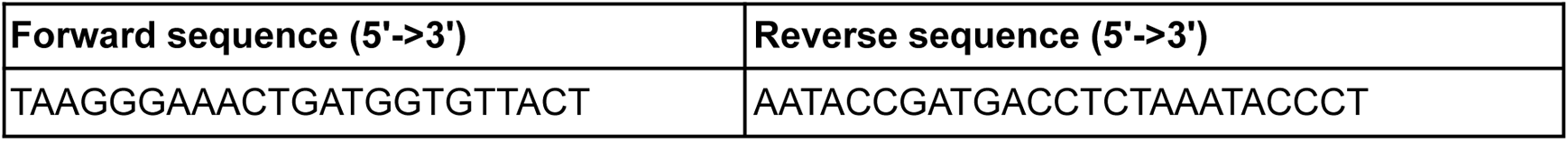
Pair of primers used for *Cdh1* promoter bisulfite PCR and Sanger sequencing. .

### Statistical analysis

Experimental data was tested for normal distribution in Prism 7 (GraphPad). Normal distributed datasets were compared with unpaired two-tailed Student’s t-tests. Comparisons of non-normal distributed datasets were performed using two-tailed Mann–Whitney U-tests. 5% significance was used. Asterisks of statistical significance (**p < 0.01, and *p < 0.05) or not significant (NS) refer to a comparison with the control, unless otherwise indicated. Embryonic, placental and blood-derived samples were allocated into experimental groups according to LPS administration. No predetermination of sample sizes was performed.

## Author contributions

DN, LA, and MRPB contributed to conceptualization, methodology, formal analysis, and interpretation of results. HMSB was responsible for animal maintenance, and ECML performed genotyping. DN carried out the investigation, collected the data, and prepared the original draft of the manuscript. DN, LA, and MRPB contributed to review and editing of the final version.

## Acknowledgement

We express our gratitude for Dr. José Carlos Farias Alves Filho for discussions on experimental designs and results. We also thank Dr. Maria Fernanda Castro-Amarante for flow cytometry experiments assistance, Dr. Simone Gomes Ferreira for µCT acquisitions and reconstruction, and the Human Genome and Stem Cell Research Center sequencing service.

